# Environmental DNA vs. Community Science: Strengths and Limitations for Urban Odonata Surveys

**DOI:** 10.1101/2024.11.26.625270

**Authors:** Rhema Uche-Dike, Ethan Tolman, Christian Benischek, Magnolia Schneider, Manpreet K. Kohli, Jonas Bush, Paul B. Frandsen, Isabella Errigo, Wendy Frankel, Kristin Gnojewski, Kim Chmura, Dick Jordan, Hannah Kittler, Michel Liao, Trinity Tobin, Cindy Su, Gracie Castillo, Ethan Derdarian, Maleah Wei, Santiago Fernandez-Jaurez, Towako Tamano, Ben Gallafent, James Jenson, Christoph A. Walser, Jessica L. Ware, Christopher D. Beatty

## Abstract

The study of insect decline remains a major frontier in insect biodiversity and conservation. Despite growing concern about accelerating rates of insect decline generally, relatively little data has been compiled about species of aquatic insects. Data is particularly lacking on the distribution of aquatic insects in urban ecosystems. Here, we compare environmental DNA (eDNA) metabarcoding and community science observation as means of monitoring Odonata within an urban system in Southwest Idaho. We show that the distribution of Odonata across this urban landscape is not uniform and that both monitoring methods have different strengths and weaknesses. We found that eDNA metabarcoding was very sensitive to the identification of genera from underrepresented families in the region, but was unable to distinguish between closely related genera, particularly from localities where eDNA could accumulate more damage. On the other hand, community science observations effectively identified the presence of genera from more speciose families but missed the presence of relatively rare species, and those that had a short flight season. These findings suggest that, in our study system, eDNA and community science are highly complementary of each other. In cases where only one method is employed for a monitoring or conservation project, care should be given to account for the biases of each approach.

## Introduction

Insect decline is a growing concern worldwide, yet little information is currently available on the extent or rate of this decline (Dicks et al., 2024; Sánchez-Bayo & Wyckhuys, 2019; Wagner et al., 2021). Aquatic insects are particularly overlooked in studies of insect decline, leaving it uncertain if their populations are experiencing similar declines to those observed for insects in terrestrial ecosystems. This gap is of particular concern given the rapid urbanization of natural habitats, which has a negative impact on aquatic macroinvertebrates (Gál et al., 2019). Odonata (dragonflies and damselflies) are of significant interest in urban ecosystems because they serve critical ecological functions—as bioindicators of environmental health and as biocontrol agents of pests like mosquitoes (Catling, 2005; Priyadarshana & Slade, 2023; Samways, 2024).

Although little is known about the distribution of Odonata in urban settings, community science, through applications such as “iNaturalist” and the collection of environmental DNA (eDNA) are two promising, non-destructive approaches. The size and unique life cycle of Odonata, involving both aquatic and terrestrial phases, make the order more readily observable by community scientists than most other aquatic insects (Dillon et al., 2022), with a wealth of data uploaded to community science-based tools such as iNaturalist. Community science has become a valuable tool in ecosystem monitoring, contributing to species data, rare species identification, and even environmental accountability, with citizen-led documentation of environmental infractions (Brooks et al., 2019; Wilson et al., 2020).

eDNA monitoring is less straightforward. This monitoring provides a non-invasive and effective means of detecting species through traces like shed cells/tissues and fecal material deposited in the environment. The approach effectively recovers biodiversity information beyond what traditional sampling methods typically reveal, particularly when a DNA metabarcoding approach is implemented (Hermans et al., 2018). Metabarcoding involves extracting DNA from bulk tissue or environmental samples, amplifying specific DNA regions (or “barcodes”), and sequencing them to identify species within a community (Deiner et al., 2015; Deiner & Altermatt,2014). eDNA metabarcoding allows for the detection of entire aquatic communities through genetic markers and captures broader ecological snapshots (Blabolil et al., 2021; Rivera et al., 2023; Yamamoto et al., 2017) that are crucial for monitoring and managing urban biodiversity.

Gaps in knowledge persist, especially regarding Odonata diversity across urban gradients and habitat types. Current limitations of eDNA methods include an inability to determine the life stage of detected organisms and a potential for missing rare species in sparse populations (Krol et al., 2019). The cost of large-scale eDNA sampling remains high, making it essential to integrate it with community science observations, which engage the public and offer extensive observational coverage. Together, these two methodologies have the opportunity to enhance biomonitoring efforts by broadening data coverage, facilitating public engagement, and strengthening conservation efforts.

Southwest Idaho serves as an ideal urban setting for Odonata monitoring using eDNA and community science. The region’s diverse aquatic habitats—rivers, ponds, wetlands, and reservoirs—support a variety of Odonata species, allowing for comparative studies of species presence across different habitat types and urban gradients. Within the region, Boise benefits from a well-established network of community scientists who are actively involved in biodiversity monitoring. This network enables comprehensive data collection across multiple water bodies, complementing eDNA data with community-led observations and fostering public involvement in urban conservation efforts. This dual approach not only strengthens the depth and breadth of urban ecosystem data but also enhances awareness and engagement around urban biodiversity.

This study aims to assess the effectiveness of integrating eDNA sampling with community science observations for monitoring Odonata in urban environments. Specifically, we evaluate the habitat-specific species presence of Odonata across urban water bodies. We compare findings from eDNA metabarcoding and direct community observations, and we propose a replicable framework for combining these techniques in urban biodiversity studies to inform conservation strategies and urban ecosystem management.

## Methods

### Environmental DNA Sampling

We selected Southwest Idaho, USA, including the cities of Boise, Meridian, Eagle, Caldwell, and McCall, as our primary sampling region (Fig. 1). This area was particularly suitable for our study due to the activity of the Boise Chapter of the Migratory Dragonfly Watch, a volunteer-led organization under the Parks and Recreation Department’s Pond Watch program. This organization had already collected a substantial baseline of migratory dragonfly data and had committed to gathering additional, comprehensive observations on all dragonfly species encountered. This robust foundation of community science data provided an excellent basis for comparing traditional observational data with environmental DNA (eDNA) monitoring methods.

**Figure 1:**
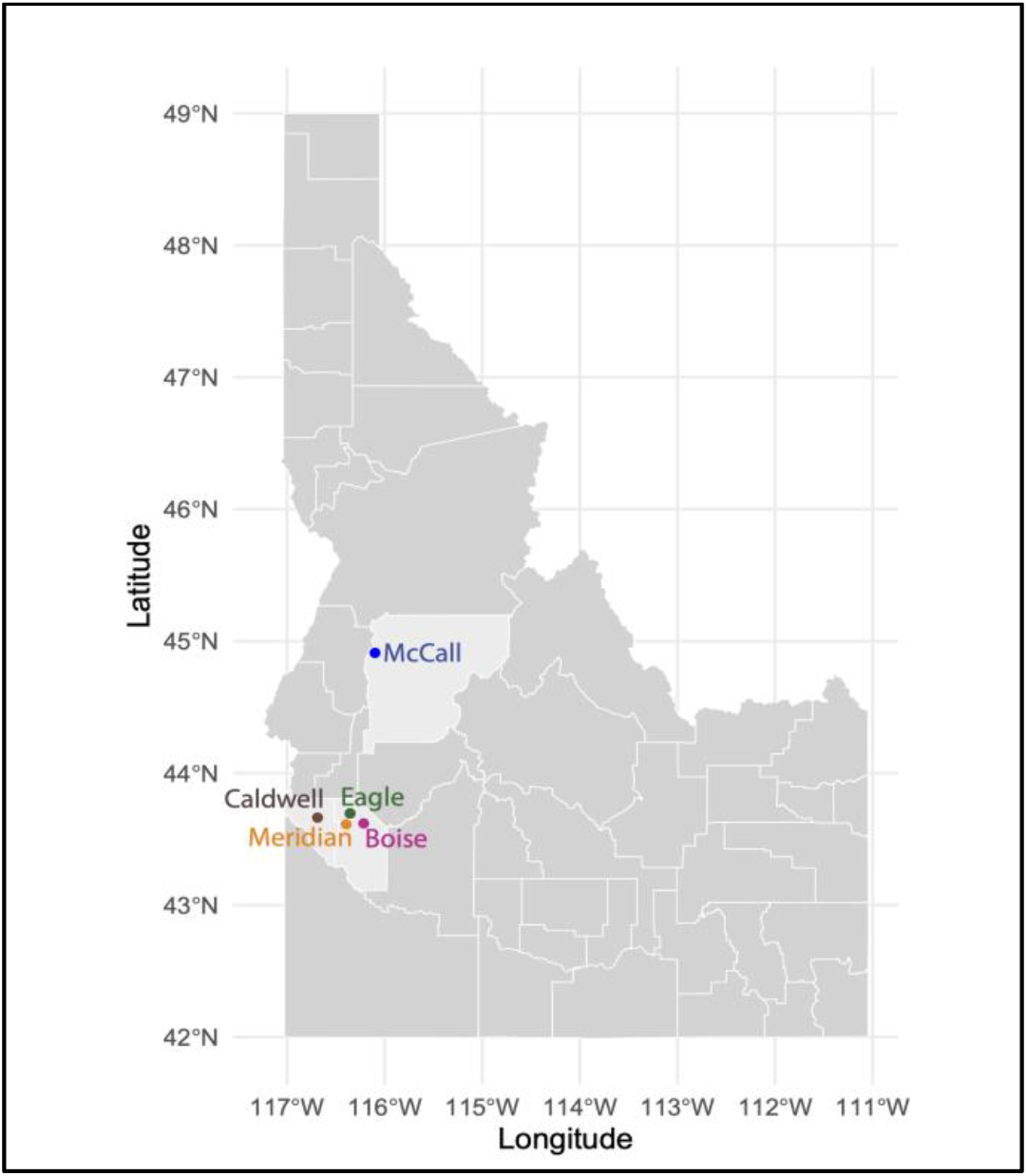
Map of Idaho, highlighting the five cities with the sampling locations. Each color indicates a different city. Light gray shade highlights counties with the sampling locations

In July 2022, we gathered water samples from 52 sites across Southwest Idaho (Supplementary Table 1), chosen for their relatively high iNaturalist density and accessibility. Our methodology largely followed previously established protocols (Pilliod et al., 2013). At each site, using VWR media/storage bottles, we collected one liter of water, which was filtered in the lab using 500 ml Thermo Scientific Nalgene filter towers and 0.45 µm filter membranes.

Because previous studies indicated that “grab-and-hold” (collecting and storing samples for later filtration) and “grab-and-filter” (immediate filtration) methods show no significant difference in DNA yield or OTU detection (Pilliod et al., 2013, 2014), filters were then stored in cryovials and preserved in a -20C freezer to maintain DNA integrity for processing. To assess the potential for laboratory contamination of our samples, we also collected tap water from the lab space where samples were filtered and filtered this water alongside field samples.

We extracted the DNA from the filters using a ZymoBIOMICS DNA miniprep kit following the manufacturer’s protocol. Given the potential levels of urea, calcium ions, and other PCR inhibitors across our study sites, we applied a targeted approach to reduce contamination. After isolating the DNA and before proceeding to PCR cycling, we performed PCR inhibitor removal, using the ZymoBIOMICS OneStep PCR inhibitor removal kit. We used two primer sets to amplify 180bp (Leese et al., 2021; Vamos et al., 2017), and 400bp (Zhou et al., 2009) regions of COI, both of which were designed specifically designed to target insects (Table 1). The forward and reverse primers adapted from Zhou et al (2009) were modified to increase amplification success for our target taxa - dragonflies and damselflies, including degenerate bases. In both primer sets we used the Apex Hot Start Taq BLUE Master Mix. For the 1709/2191 primer set (Table 1), thermal cycling conditions included an initial denaturation at 95°C for 15 minutes, followed by 35 cycles of denaturation at 95°C for 30 seconds, annealing at 51°C for 30 seconds, and extension at 72°C for 2 minutes and 30 seconds. A final elongation step was carried out at 68°C for 10 minutes, and samples were held at 4°C. For the fwhF2/EPTDr2n primer set, the protocol was similar, with the annealing temperature adjusted to 50.1°C and identical extension and final elongation steps.

**Table 1:**
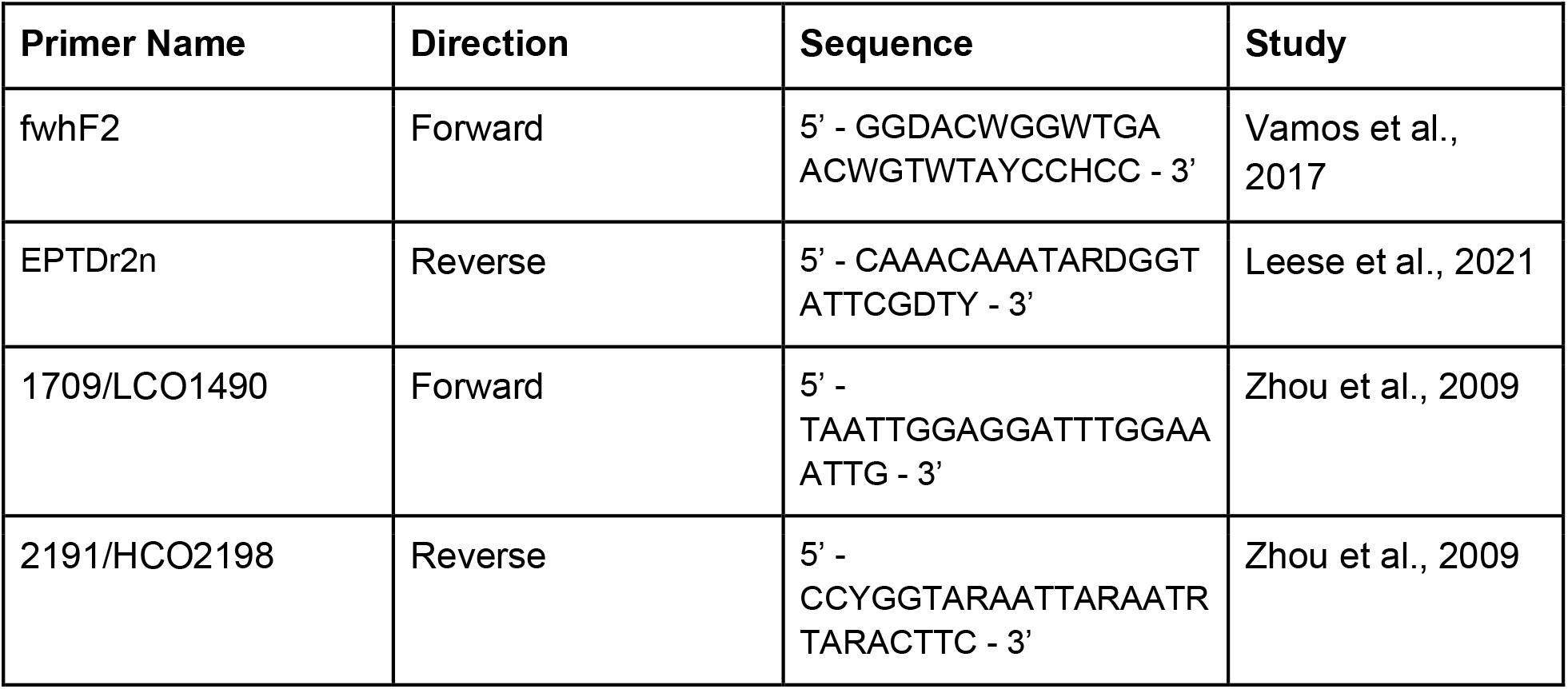
List of COI primers used in this study, including modified sequences containing degenerate bases, accounting for variation in the target sequence.

To increase the reproducibility of our methodology, we sequenced the amplicons on the Oxford Nanopore MK1C, a relatively affordable DNA sequencing machine. PCR amplicons for each site were pooled, and barcodes were attached using the ONT Native Barcoding Kit 96 V 10 chemistry for library preparation. To reduce the impact of off-target amplification we then pooled all barcoded samples, ran the pooled samples through a .5% agarose gel, and isolated loci of interest, based on size, using the ThermoFisher GeneJet gel extraction kit. We then proceeded with library preparation according to the manufacturer’s protocols, and sequenced the library on a R10.4.1 flow cell, with “high accuracy” base-calling enabled, which reduces the error rate below 10 percent (Sanderson et al., 2023).

#### Bioinformatic analysis

We processed sequencing data by clustering reads at 98% similarity using CD-HIT v4.8.1 (Fu et al., 2012), which reduces redundancy and groups sequences into molecular operational taxonomic units (MOTUs) while maintaining taxonomic resolution. This threshold is commonly used in eDNA studies targeting metazoans to balance computational efficiency and genus-level resolution. We used BlastN (Camacho et al., 2009) to compare the clustered sequences against the GenBank nucleotide database. An e-value threshold of 1e-40 and a percent identity cutoff of 70% were applied to filter high-confidence matches while accounting for genetic variability in mitochondrial markers (Ratnasingham & Hebert, 2007).

To refine our dataset, we retained only sequences with significant hits to Eumetazoa, focusing on target taxa while excluding potential contaminants such as prokaryotes. Clustered OTUs were then identified using Boldigger v2.1.1 (Buchner & Leese, 2020), which integrates data within the Barcode of Life Data Systems (BOLD). Boldigger enables taxonomic assignment through BOLD’s extensive reference library, which is widely used in biodiversity research.

### Community Scientist Recruitment

For immediate comparison with eDNA data in the same field season, we pulled all iNaturalist observations for Odonata at our 52 study sites in 2022 and recorded the presence/absence of each genus. To establish a longer-term community baseline for Southwest Idaho we worked with “Pondwatch” volunteers at the City of Boise who intensively observed a subset of eight of the 52 sites (chosen for accessibility) for the field seasons of 2022, 2023, and 2024 (Supplementary Table 2). In 2022 volunteers were instructed to also upload their observations to iNaturalist to supplement passive community science observations. These volunteers were first trained in 2019; when Boise Parks and Recreation staff provided a classroom to observe migratory species, and since 2022, the program has expanded by encouraging volunteers to record all observed Odonata, with yearly training for species identification.

## Results

In Idaho, Odonata DNA was detected at 27 of the 52 sites sampled (52%), at 16 (59%) of these sites Odonata were also recorded on iNaturalist (Fig. 2). Three of the families observed on iNaturalist–Cordulegastridae, Lestidae, and Calopterygidae–were not detected through eDNA (Fig. 2). Gomphidae and Corduliidae were the two most commonly identified families in the eDNA data, despite the fact that, in the iNaturalist observations, Gomphidae were only observed at three sites, and Corduliidae were observed at one (supplementary table 1). Libellulidae were identified from eDNA at ten sites, three of which were in locations where they were also identified on iNaturalist (supplementary tables 1,3, Fig. 3). Aeshnidae were only identified at two sites, one of which was a site where Aeshnidae were identified on iNaturalist (supplementary tables 1,3). Coenagrionidae were identified at five sites, two of which were sites where Coenagrionidae were uploaded to iNaturalist (supplementary tables 1,3, Fig. 2). Of the OTUs identified to a genus level, *Phanogomphus* (*Phanogomphus kurilis*), *Erythemis* (*Erythemis collocata*), and *Pachydiplax* (*Pachydiplax longipennis*) were from genera containing only one species and can be presumed to be species-level OTUs.

**Figure 2:**
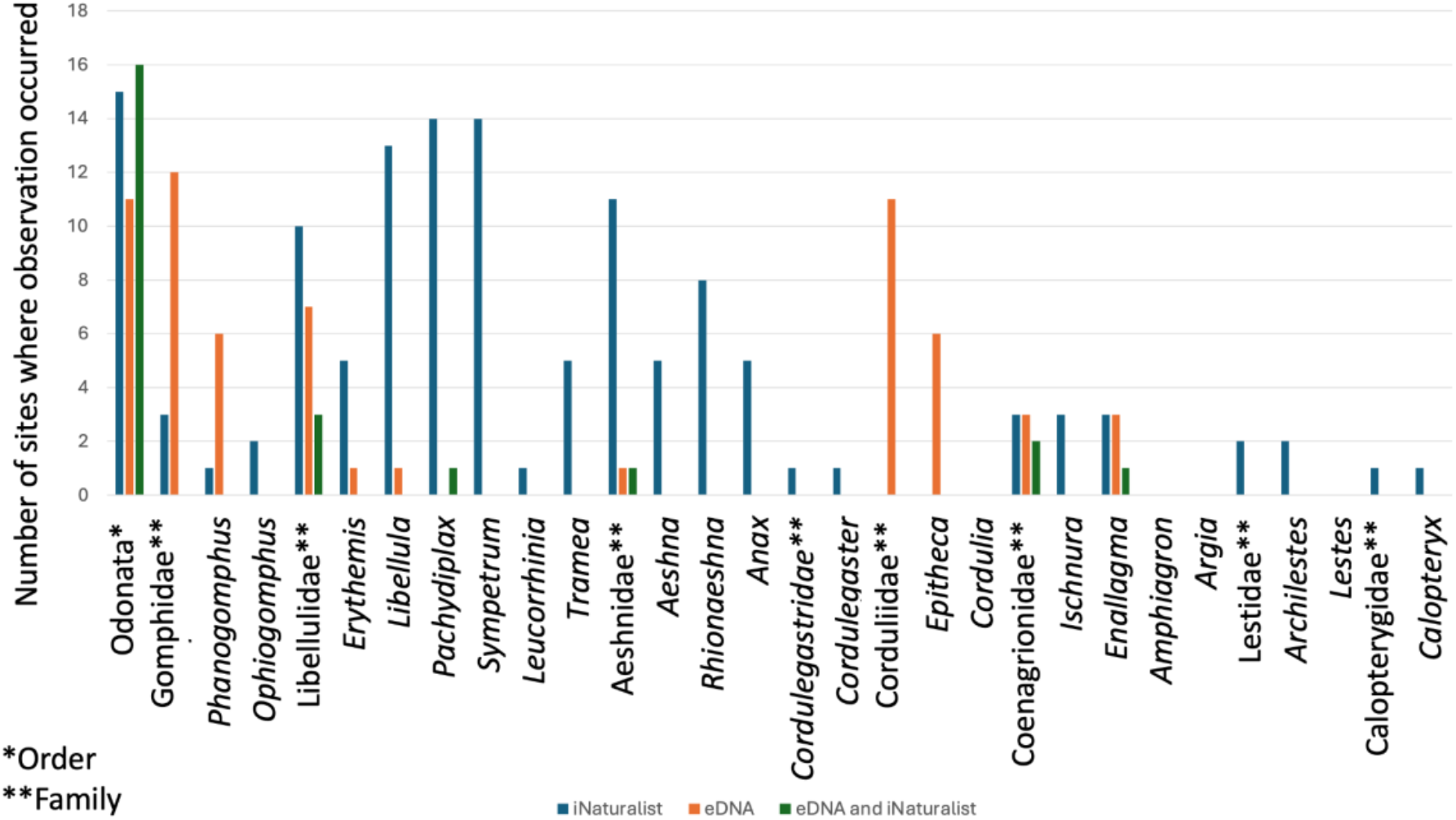
Presence of Odonata from eDNA and iNaturalist sampling. Histogram displaying the taxa richness; where each genus or family was identified through eDNA and/or iNaturalist observations. Pondwatch observations were uploaded to iNaturalist and were included in this analysis.

**Figure 3:**
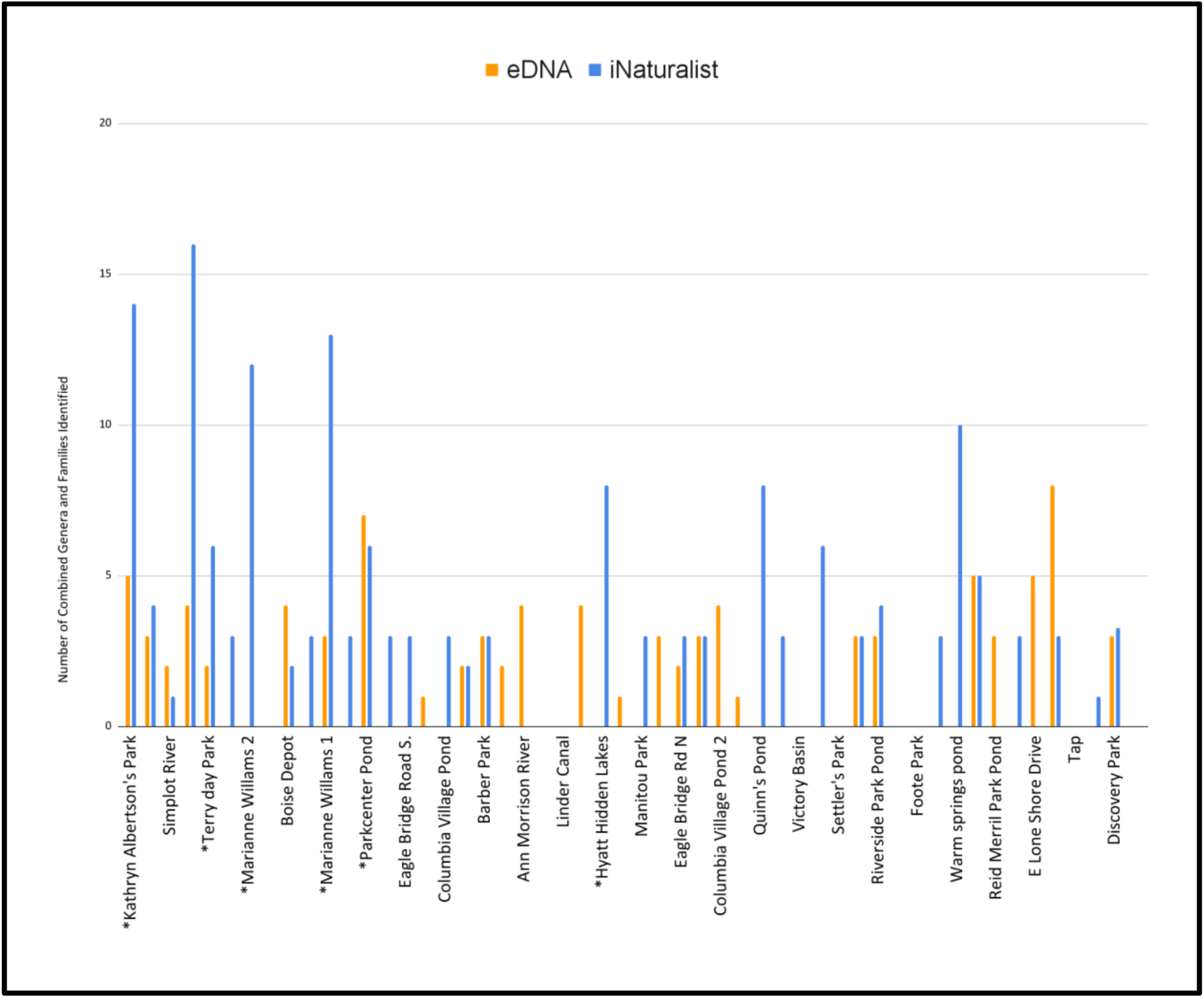
Taxa richness per site from iNaturalist observations (blue) and OTUs identified from eDNA (orange). Sites include observations from iNaturalist data and Pondwatch observation data. Sites with asterisks included observations from Pondwatch volunteers. An average of 3.25 genera were observed per site.

### Community science

In our 52 sites, we pulled the presence of eight families and 21 genera (Supplementary Table 3) from iNaturalist. This included 5 families of Anisoptera (Aeshnidae, Cordulegastridae, Corduliidae, Gomphidae, and Libellulidae), and 3 families of Zygoptera (Coenagrionidae, Lestidae, and Calopterygidae). Odonata was observed at 30 of the 52 sites where water samples were collected (Supplementary Table 3). Over the more intensive three-year monitoring of a subset of eight sites, city volunteers observed 12 genera at their monitored localities, including one additional family, Macromiidae (Fig.4), not recorded by other sampling efforts.

## Discussion

This study synthesizes the distribution of Odonata in the Boise metropolitan area. We found a high degree of discordance between our two survey methods (Fig. 2, Supplementary table 1), likely driven by the unique biology and life histories making up the Odonata assemblages of Southwest Idaho, and the advantages of both community science and eDNA. By combining results from eDNA, iNaturalist, and pondwatch observations, we also identified Kathryn Albertson’s Pond, Veterans Pond, and Marianne Williams Park as localities of high odonate richness in our study area (Supplementary Table 1, Fig. 3).

### Sensitivities of two biomonitoring methods

*Species underrepresented in eDNA sampling compared to community science*

A large proportion of the molecular OTUs could not be identified past the family level (Supplementary table 1). This was especially true of Libellulidae, a species-rich family, most of which can be considered to be generalists in western North America (Paulson, 2009). This was, by far, the most commonly observed family by community scientists (Supplementary table 2), but highly common genera such as *Pachydiplax* and *Sympetrum* rarely showed up in eDNA (Fig. 2). There are several possible explanations for this. For example, many Libellulidae are relatively recently diverged (Goodman et al., 2023), and COI may not be a reliable identifier to the species, or even genus level, as is the case in small numbers of birds, fishes (Ward, 2009),

and flies (Lin et al., 2015). Additionally, Libellulidae can live in highly polluted urban water bodies where other species cannot be found, and their gene family evolution suggests shared pressures to limit and repair DNA damage (Tolman et al., 2023). Chemical compounds from pollutants can alter water chemistry and rapidly degrade environmental DNA (Joseph et al., 2022), potentially contributing to the reduced persistence of eDNA in the habitats where Libellulidae are most likely to be found. These biological difficulties could be further compounded by the sequencing error in the Oxford Nanopore Technologies library used (∼10%). Newer sequencing chemistries and base-calling algorithms have already improved the error rate of Oxford Nanopore sequences, potentially alleviating this source of error in the future.

#### Species overrepresented in eDNA compared to community science

The Pacific Clubtail (*Phanogomphus kurilis*), the only species of *Phanogomphus* found in our study area, was identified at six different localities using eDNA (Kathryn Albertson Park, Parkcenter Pond, Crab Shack, Columbia Village Pond 2, Magnolia Park, and E Lone Shore Drive) but was only observed in one location in community science efforts (Fig. 4). *P. kurilis* typically inhabit rocky lotic water as larvae, before emerging and surviving for two to three weeks as adults (Paulson, 2009). This OTU could simply be much easier to identify bioinformatically, as it is the only member of its genus known to inhabit Southwest Idaho.

**Figure 4:**
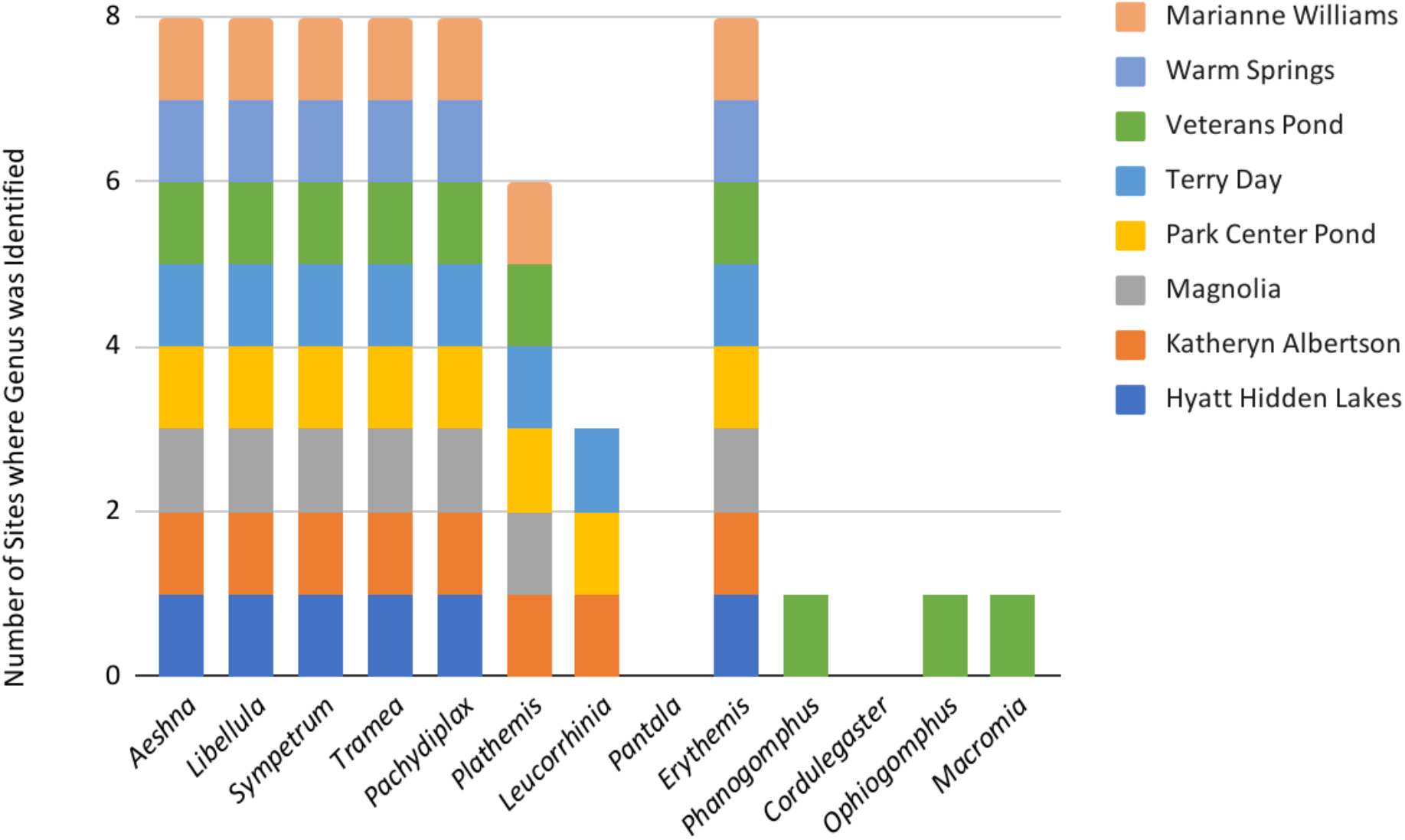
Composition of “Pondwatch” Observations. Taxa richness per site from Pondwatch observations by genus. Sampling was carried out from 2022-2024. Marianne Williams Ponds 1 and 2 are combined in this dataset.

Intriguingly, eDNA was consistently detected upstream of its community science locality, and even in lentic habitats (such as Columbia Village Pond 2), despite upstream-related challenges like water flow and spatial heterogeneity highlighted in previous studies (Sansom et al., 2024; Wood et al., 2021). While the community science locality no doubt represents a breeding population, we hypothesize that a second breeding locality exists upstream of our sampled localities. Odonata generally (R. B. DuBois, 2020), and Gomphidae specifically (B. DuBois & Pratt, 2017) are known to drift downstream as larvae, as could their DNA (Deiner & Altermatt, 2014). *P. kurilis* is a lotic species, and the drift of its eDNA, in particular, is a plausible explanation for their identification in lentic habitats (Supplementary Table 1), given that the lentic habitats sampled all receive input from streams and rivers.

OTUs from eDNA sequencing were assigned to the family Corduliidae at 11 sites (Supplementary table 1), but this family was not observed by community scientists during our study period at any of our 52 sites in 2022, nor in the subset of eight sites observed by Pondwatch volunteers through 2024 (Fig. 4). *Somatochlora* and *Cordulia* are the most abundant genera of this family in Idaho (Paulson, 2009), and both have been observed throughout Southwest Idaho more broadly, outside of our temporal study period. The behavior of *Somatochlora*, which is classified as a flier, and rarely perches (Paulson, 2009), could make it difficult to capture in passive observations. *Cordulia* has a flight season in late May (Paulson, 2009), earlier than many other species. This could partially explain its absence from community science observations, which tend to occur later in Summer. While the genus *Epitheca* was identified in our eDNA sampling at six sites (Fig. 3), it has not previously been identified in Southern Idaho. These are highly noticeable odonates, with spotted wings, and we find it unlikely that these would have been overlooked by iNaturalist observers. This could represent a range extension, but it is possible that the signal of *Epitheca* is an artifact. Further research will be needed to determine how both life history and our eDNA pipeline impact the detection of species.

### Conservation and monitoring implications

Our eDNA pipeline excelled in identifying more “rare” species that do not have any close relatives in the study region, while community science observations capture more common genera, and can more readily distinguish between closely related species. As Odonata are important indicators for the health of aquatic ecosystems (Catling, 2005; Kietzka et al., 2022) this finding has practical applications for how Odonata monitoring could be used in practice. If the relative species richness of Odonata is to be used as a surrogate for aquatic biodiversity, community science observations would indeed be a very useful metric. However, if the presence of a relatively rare taxon is being used as a biodiversity surrogate, our eDNA pipeline may be more sensitive.

### Odonata Distribution in Urban Southwest Idaho

Kathryn Albertson’s Pond had a particularly high number of Odonata observations in our iNaturalist survey with 66 of the 383 (∼17%) total sightings and 14 observed genera and families recorded (compared to the 3.235 taxa that were identified at each pond site on average).

Kathryn Albertson Park was intentionally designed to support urban biodiversity (“Kathryn Albertson Park,” n.d.), with a dense assemblage of native plant life, potentially making it a feeding ground for Odonata adults (which can feed in different habitats than reproduction and oviposition (Paulson, 2009)), while larvae may not be abundant in the water. This park also harbored unique taxa and was the only locality where the genus *Archilestes* was found using both iNaturalist and eDNA data (Supplementary tables 1,3). The intentional planting of native aquatic plants may have created more of the habitat that *Archilestes* needs to survive as a nymph (Paulson, 2009).

Veterans Park Pond was similarly noted for its genus diversity, including 16 genera and families identified using iNaturalist, and accounting for 15.7% of the total Odonata observations (Fig. 3). This was the only site where the Pacific Clubtail (*Phanogomphus*), Sinuous Snaketail (*Ophiogomphus*), and Western River Cruiser (*Macromia*) were observed with either community science approach (Fig. 3, Fig. 4). This pond is noted for its relatively large surface area (17.6 acres) in comparison to other lentic habitats considered in this study, perhaps providing more micro-habitats.

A further 29.69% of all iNaturalist observations occurred in Marianne Williams Park, a 72-acre park with multiple ponds and storm drainages. Such habitat diversity has been shown to support higher aquatic insect composition (Santi et al., 2010).

While the number of community science observations at each site could be a reflection of uneven sampling effort, it may not be the case. The eight “Pond Watch Sites”, which would have had dedicated volunteers making iNaturalist observations, did not have the same number of observations. Additionally, other sites near the city center (such as Ann Morrison and Julia Davis Parks) did not have a high number of Odonata observations (Supplementary Table 3).

However, the inability to track sampling effort is a major drawback of passive sampling like iNaturalist. While the aforementioned sites are variable in their topography, they are all in close proximity to the Boise River. While we are not aware whether odonates are using the river as a corridor through the city, this could explain the higher diversity at sites adjacent to it, although this has not been studied in Odonata. We do not have the data to test this hypothesis here, but this is an exciting area of future study.

## Conclusions

Our combined datasets reveal that community science can provide extensive and valuable distribution data, while eDNA metabarcoding using the relatively accessible Oxford Nanopore MinION is an effective tool for detecting unique Odonata genera within a community assembly, though it has limitations in distinguishing between closely related taxa. Furthermore, we raise the possibility that both the proximity to the river and pond design can have a dramatic influence on Odonata assemblages in urban habitats, impacting species diversity and distribution patterns.

Additionally, for charismatic groups such as Odonata, in which the community readily engages on community sciences platforms, our study shows passive community science surveys and documentation can provide very informative data at a fraction of the cost. This is particularly useful for regions in the global south where there are limited resources for methods like eDNA, in addition to the regional environmental challenges that eDNA sampling poses.

## References

Blabolil, P., Harper, L. R., Říčanová, Š., Sellers, G., Di Muri, C., Jůza, T., Vašek, M., Sajdlová, Z., Rychtecký, P., Znachor, P., Hejzlar, J., Peterka, J., & Hänfling, B. (2021). Environmental DNA metabarcoding uncovers environmental correlates of fish communities in spatially heterogeneous freshwater habitats. Ecological Indicators, 126, 107698. 10.1016/j.ecolind.2021.107698

Brooks, S. J., Fitch, B., Davy-Bowker, J., & Codesal, S. A. (2019). Anglers’ Riverfly Monitoring Initiative (ARMI): A UK-wide citizen science project for water quality assessment. Freshwater Science, 38(2), 270–280. 10.1086/703397

Catling, P. M. (2005). A Potential for the Use of Dragonfly (Odonata) Diversity as a Bioindicator of the Efficiency of Sewage Lagoons. The Canadian Field-Naturalist, 119(2), 233. 10.22621/cfn.v119i2.111

Deiner, K., & Altermatt, F. (2014). Transport Distance of Invertebrate Environmental DNA in a Natural River. PLOS ONE, 9(2), e88786. 10.1371/journal.pone.0088786

Deiner, K., Walser, J.-C., Mächler, E., & Altermatt, F. (2015). Choice of capture and extraction methods affect detection of freshwater biodiversity from environmental DNA. Biological Conservation, 183, 53–63. 10.1016/j.biocon.2014.11.018

Dicks, L. V., Grames, E. M., Bowler, D. E., & Isaac, N. J. B. (2024). Insect declines – an overview of current knowledge on the status of the world’s insects. In J. S. Pryke, M. J. Samways, T. R. New, P. Cardoso, & R. Gaigher, Routledge Handbook of Insect Conservation (1st ed., pp. 77–91). Routledge. 10.4324/9781003285793-8

Dillon, A., Simaika, J., Clausnitzer, V., Thompson, A., White, E., Montes-Fontalvo, J., Goforth, C., & Khelifa, R. (2022). Bridging people and nature through Odonata. In A. Cordoba-Aguilar, C. Beatty, & J. Bried (Eds.), Dragonflies and Damselflies: Model Organisms for Ecological and Evolutionary Research (p. 0). Oxford University Press. 10.1093/oso/9780192898623.003.0029

DuBois, B., & Pratt, D. (2017). Species and life stages of Odonata nymphs sampled with large drift nets in two Wisconsin rivers. The Great Lakes Entomologist, 50(1). 10.22543/0090-0222.1000

DuBois, R. B. (2020). Odonata drift: A reassessment. International Journal of Odonatology, 23(4), 381–396. 10.1080/13887890.2020.1818639

Gál, B., Szivák, I., Heino, J., & Schmera, D. (2019). The effect of urbanization on freshwater macroinvertebrates – Knowledge gaps and future research directions. Ecological Indicators, 104, 357–364. 10.1016/j.ecolind.2019.05.012

Goodman, A., Tolman, E., Uche-Dike, R., Abbott, J., Breinholt, J. W., Bybee, S., Frandsen, P. B., Gosnell, J. S., Guralnick, R., Kalkman, V. J., Kohli, M., Lontchi, J. F., Lupiyaningdyah, P., Newton, L., & Ware, J. L. (2023). Assessment of targeted enrichment locus capture across time and museums using odonate specimens. Insect Systematics and Diversity, 7(3), 5. 10.1093/isd/ixad011

Hermans, S. M., Buckley, H. L., & Lear, G. (2018). Optimal extraction methods for the simultaneous analysis of DNA from diverse organisms and sample types. Molecular Ecology Resources, 18(3), 557–569. 10.1111/1755-0998.12762

Joseph, C., Faiq, M. E., Li, Z., & Chen, G. (2022). Persistence and degradation dynamics of eDNA affected by environmental factors in aquatic ecosystems. Hydrobiologia, 849(19), 4119–4133. 10.1007/s10750-022-04959-w

Kathryn Albertson Park. (n.d.). Biology - The College of Arts and Sciences. Retrieved October 19, 2024, from https://www.boisestate.edu/biology/about/about-our-department__trashed/environment/kathryn-albertson-park/

Kietzka, G. J., Deacon, C., & Patten, M. A. (2022). Odonata as surrogates of biodiversity. In A. Cordoba-Aguilar, C. Beatty, & J. Bried (Eds.), Dragonflies and Damselflies: Model Organisms for Ecological and Evolutionary Research (p. 0). Oxford University Press. 10.1093/oso/9780192898623.003.0025

Krol, L., Van der Hoorn, B., Gorsich, E. E., Trimbos, K., Bodegom, P.M. van, & Schrama, M. (2019). How Does eDNA Compare to Traditional Trapping? Detecting Mosquito Communities in South-African Freshwater Ponds. Frontiers in Ecology and Evolution, 7. https://www.frontiersin.org/article/10.3389/fevo.2019.00260

Leese, F., Sander, M., Buchner, D., Elbrecht, V., Haase, P., & Zizka, V. M. A. (2021). Improved freshwater macroinvertebrate detection from environmental DNA through minimized nontarget amplification. Environmental DNA, 3(1), 261–276. 10.1002/edn3.177

Lin, X., Stur, E., & Ekrem, T. (2015). Exploring Genetic Divergence in a Species-Rich Insect Genus Using 2790 DNA Barcodes. PLOS ONE, 10(9), e0138993. 10.1371/journal.pone.0138993

Paulson, D. (2009). Dragonflies and Damselflies of the West. Princeton University Press.

Pilliod, D. S., Goldberg, C. S., Arkle, R. S., & Waits, L. P. (2013). Estimating occupancy and abundance of stream amphibians using environmental DNA from filtered water samples. Canadian Journal of Fisheries and Aquatic Sciences, 70(8), 1123–1130. 10.1139/cjfas-2013-0047

Pilliod, D. S., Goldberg, C. S., Arkle, R. S., & Waits, L. P. (2014). Factors influencing detection of eDNA from a stream-dwelling amphibian. Molecular Ecology Resources, 14(1), 109– 116. 10.1111/1755-0998.12159

Priyadarshana, T. S., & Slade, E. M. (2023). A meta-analysis reveals that dragonflies and damselflies can provide effective biological control of mosquitoes. Journal of Animal Ecology, 92(8), 1589–1600. 10.1111/1365-2656.13965

Ratnasingham, S., & Hebert, P. D. N. (2007). bold: The Barcode of Life Data System (http://www.barcodinglife.org). Molecular Ecology Notes, 7(3), p355–364. 10.1111/j.1471-8286.2007.01678.x

Rivera, S. F., Vasselon, V., Bouchez, A., & Rimet, F. (2023). eDNA metabarcoding from aquatic biofilms allows studying spatial and temporal fluctuations of fish communities from Lake Geneva. Environmental DNA, 5(3), 570–581. 10.1002/edn3.413

Samways, M. J. (2024). Freshwater Assessment and Monitoring Using Dragonflies. In Conservation of Dragonflies (pp. 331–438). 10.1079/9781789248395.0007

Sánchez-Bayo, F., & Wyckhuys, K. A. G. (2019). Worldwide decline of the entomofauna: A review of its drivers. Biological Conservation, 232, 8–27. 10.1016/j.biocon.2019.01.020

Sanderson, N. D., Kapel, N., Rodger, G., Webster, H., Lipworth, S., Street, T. L., Peto, T., Crook, D., & Stoesser, N. (2023). Comparison of R9.4.1/Kit10 and R10/Kit12 Oxford Nanopore flowcells and chemistries in bacterial genome reconstruction. Microbial Genomics, 9(1), mgen000910. 10.1099/mgen.0.000910

Sansom, B. J., Ruiz-Ramos, D. V., Thompson, N. L., Roberts, M. O., Taylor, Z. A., Ortiz, K., Jones, J. W., Richter, C. A., & Klymus, K. E. (2024). Detection and transport of environmental DNA from two federally endangered mussels. PLOS ONE, 19(10), e0304323. 10.1371/journal.pone.0304323

Tolman, E. R., Beatty, C. D., Frandsen, P. B., Bush, J., Bruchim, O. R., Driever, E. S., Harding, K. M., Jordan, D., Kohli, M. K., Park, J., Park, S., Reyes, K., Rosari, M., Ryu, J. L., Wade, V., & Ware, J. L. (2023). Newly Sequenced Genomes Reveal Patterns of Gene Family Expansion in select Dragonflies (Odonata: Anisoptera) (p. 2023.12.11.569651). bioRxiv. 10.1101/2023.12.11.569651

Vamos, E., Elbrecht, V., & Leese, F. (2017). Short COI markers for freshwater macroinvertebrate metabarcoding. Metabarcoding and Metagenomics, 1, e14625. 10.3897/mbmg.1.14625

Wagner, D. L., Grames, E. M., Forister, M. L., Berenbaum, M. R., & Stopak, D. (2021). Insect decline in the Anthropocene: Death by a thousand cuts. Proceedings of the National Academy of Sciences, 118(2). 10.1073/pnas.2023989118

Ward, R. D. (2009). DNA barcode divergence among species and genera of birds and fishes. Molecular Ecology Resources, 9(4), 1077–1085. 10.1111/j.1755-0998.2009.02541.x

Wilson, J. S., Pan, A. D., General, D. E. M., & Koch, J. B. (2020). More eyes on the prize: An observation of a very rare, threatened species of Philippine Bumble bee, Bombus irisanensis, on iNaturalist and the importance of citizen science in conservation biology. Journal of Insect Conservation, 24(4), 727–729. 10.1007/s10841-020-00233-3

Wood, Z. T., Lacoursière-Roussel, A., LeBlanc, F., Trudel, M., Kinnison, M. T., Garry McBrine, C., Pavey, S. A., & Gagné, N. (2021). Spatial Heterogeneity of eDNA Transport Improves Stream Assessment of Threatened Salmon Presence, Abundance, and Location. Frontiers in Ecology and Evolution, 9. 10.3389/fevo.2021.650717

Yamamoto, S., Masuda, R., Sato, Y., Sado, T., Araki, H., Kondoh, M., Minamoto, T., & Miya, M. (2017). Environmental DNA metabarcoding reveals local fish communities in a species-rich coastal sea. Scientific Reports, 7(1), 40368. 10.1038/srep40368

Zhou, X., Adamowicz, S. J., Jacobus, L. M., DeWalt, R. E., & Hebert, P. D. (2009). Towards a comprehensive barcode library for arctic life—Ephemeroptera, Plecoptera, and Trichoptera of Churchill, Manitoba, Canada. Frontiers in Zoology, 6(1), 30. 10.1186/1742-9994-6-30

