## Supplementary Table 2 for "Environmental DNA vs. Community Science: Strengths and Limitations for Urban Odonata Surveys"

| **Pond Name** | **Acreage** | **Vegetation and Pond Description** | **Winter Conditions** | **Boise River Adjacent** | **Fishing Permitted** | **Fish Stocking** | **Other Fish Present** |
| --- | --- | --- | --- | --- | --- | --- | --- |
| **Kathryn Albertson’s Park** | Unknown (several large ponds) | Several shallow ponds with abundant overhanging trees, emergent vegetation, and lilypads. | Shallow water in some sections, dry in others, water level drops significantly | Yes | No | No | Carp and other non-native species |
| **Hyatt Hidden Lakes Reserve** | Unknown (several large ponds) | Several ponds ranging from shallow to deep with abundant emergent vegetation including cattails, rushes, and willows.  Several islands with vegetation that are inaccessible to the public. | Water level drops significantly, but ponds still wet | No | No | No | Several species.  Many dumped by public. |
| [**Marianne Williams Park,**](https://idfg.idaho.gov/ifwis/fishingplanner/water/1161411435719) **Pond 1*** | 2.6 acres | Large pond with aerators.  Abundant emergent vegetation including cattails, rushes, willow, and mature cottonwood trees.  Gentle gradient from deep center portion of pond to pond edge. | Full | Yes | Yes | Rainbow trout | All observed in 2016: Bluegill Lepomis macrochirus, Largemouth Bass Micropterus salmoides, Largescale Sucker Catostomus macrocheilus, Pumpkinseed Lepomis gibbosus, Rainbow Trout Oncorhynchus mykiss |
| [**Marianne Williams Park,**](https://idfg.idaho.gov/ifwis/fishingplanner/water/1161489435757) **Pond 2*** | 3.6 acres | Large pond with aerators. Abundant emergent vegetation including cattails, rushes, willow, and mature cottonwood trees. Gentle gradient from deep center portion of pond to pond edge. | Full | Yes | Yes | Bluegill, Largemouth  bass | [Bluegill / Pumpkinseed / Sunfish(Lepomis)](https://idfg.idaho.gov/species/taxa/7257) |
| [**Parkcenter P**](https://idfg.idaho.gov/ifwis/fishingplanner/water/1161823435980)**ond*** | 8 acres | Abundant cattails and rushes surrounding entirety of pond.  Mature trees nearby and overhanging pond in some areas. | Full | No, but fairly close.  Adjacent to Logger's Creek. | Yes | Rainbow trout | [Bluegill Lepomis macrochirus observed in 2014, Brown Bullhead, Common Carp, Largemouth Bass, Largescale Sucker, Oriental Weatherfish, Pumpkinseed, Rainbow Trout, Yellow Perch](https://idfg.idaho.gov/species/taxa/16589) |
| [**Terry Day Park**](https://idfg.idaho.gov/ifwis/fishingplanner/water/1162065435914) | .7 acres | Located in a neighborhood park.  Abundant cattails and rushes surrounding entirety of pond.  Mature trees nearby and overhanging pond in some areas. | Full | No | Yes | Bluegill, Largemouth bass, Pumpkinseed | Bluegill, Largemouth Bass, Smallmouth Bass |
| **Warm Springs Pond** |  | Abundant cattails, rushes, willows, and mature trees surround entire pond area.  Shallow pond fed by irrigation water. | Drained, small, shallow remnant pond in main pond area | Yes | No | No | No fish known to be present |
| [**Veteran's Pond**](https://idfg.idaho.gov/ifwis/fishingplanner/water/1162375436324)* | 17.6 acres | Deep, cold pond.  Lacks significant emergent vegetation in many areas.  Several mature cottonwoods overhanging, but lots of exposed areas. | Full | Yes | Yes | Rainbow Trou | [Bluegill / Pumpkinseed / Sunfish(Lepomis), Largemouth Bass, Rainbow Trout](https://idfg.idaho.gov/species/taxa/7257) |
| [**Quinn’s**](https://idfg.idaho.gov/ifwis/fishingplanner/water/1162321436232) **Pond** | 23.4 | Remnant gravel pit, minimal vegetation around most sides of the pond, some cattails on the north side, mature trees (Cottonwoods, Black locust, Siberian elm), but lots of exposure.  Few perching areas adjacent to pond.  Pollinator garden to south, greenbelt to west and apartments with turf to the east.  Pond is relatively deep and near intake from Boise River. | Full | Yes | Yes | Channel catfish, Brown trout, Rainbow trout, Steelhead | Bluegill / Pumpkinseed / Sunfish (Lepomis) Largemouth Bass (Micropterus salmoides) Yellow Perch (Perca flavescens) Catfish (Ictalurus furcatus) Rainbow Trout (Oncorhynchus mykiss) |
| [**Magnolia Pond**](https://idfg.idaho.gov/ifwis/fishingplanner/water/1162922436812) | .3 acres | Located in a neighborhood park.  Some emergent vegetation including cattails, willows, and rushes.  Trees are still relatively small. | Full | No | Yes | Bluegill, Largemouth  bass | [Largemouth Bass (Micropterus salmoides), Bluegill](https://idfg.idaho.gov/species/taxa/16488) |
| **Julia Davis Park Pond** |  | Shallow, muddy pond. | Drained | Yes | No | No |  |
| *Supplementary Table 2: Descriptions of Pondwatch monitoring sites* | | | | | | | |
